# BinChecker: a new algorithm for quality assessment of microbial draft genomes

**DOI:** 10.1101/2021.10.01.462745

**Authors:** Heiner Klingenberg, Peter Meinicke

## Abstract

In the reconstruction of microbial genomes from metagenomic sequence data, the estimation of the final completeness and possible contamination is crucial for quality control. In metagenomics candidate genomes are usually obtained from a metagenome assembly and a subsequent binning of the assembled contigs. BinChecker provides a novel approach to quality assessment that is based on a fast protein domain search and a clustering approach for identification of marker domain (“feature”) sets. The feature sets that are used for estimation are not pre-computed for a given database of reference genomes, but are individually found for each bin by adaptive clustering and feature selection. In particular, the adaptivity facilitates the creation and extension of the underlying database, which just requires to add protein feature profiles of reference genomes. Tests with simulated bins indicate that the prediction accuracy of BinChecker meets the current state of the art while providing significant advantages in terms of speed and flexibility.

## 1 Introduction

The number of genomes obtained from metagenome and single cell sequencing data rapidly increases. However, the variation in quality of the resulting draft genomes is considerable [1]. For all subsequent analyses that make use of these data, information about the completeness and purity of these genomes is essential. Therefore, an important aspect is the quality assessment of the new candidate genomes in terms of completeness and contamination estimates: how much of the original genome is actually present and how much of the genomic material possibly stems from other organisms?

The standard for measuring completeness and contamination of bacterial and archaeal genomes is to count the occurrences of universal single copy marker genes in the candidate genomes. However, this common approach has some inherent limitations [2]. The first problem is that these marker genes only represent a small part of the entire genome. Universal single copy gene (SCG) sets for archaea and bacteria typically contain about 50 genes which can only cover a few percent of an average size microbial genome. In real applications the situation may even be worse because SCG appear to be clustered on genomes and therefore a bin might correspond to a biased sample. The second, closely related problem is that it is difficult if not impossible to distinguish between changes in completeness and contamination. In fact, for a mixture of two incomplete genomes, a high completeness with almost no contamination may be predicted as long as the marker genes from the two organisms complement each other.

CheckM tries to overcome these problems by using subsets of lineage-specific protein domains from the PFAM database [3] as marker sets. If a candidate draft genome can be assigned to one of the taxonomic groups represented in CheckM, the tool uses a specific marker set which can comprise more than thousand protein domains, depending on the taxonomic level and the number of reference genomes in the assigned group. The taxonomic assignment in CheckM is based on phylogenetic analysis of ribosomal proteins. This can considerably improve the completeness and contamination estimates in cases where closely related genomes are present in the CheckM database. If an assignment to a more specific group is not possible CheckM automatically uses a universal marker set.

At the heart of the CheckM approach is the identification of lineage-specific marker sets, which is the key to enlarge the statistical basis for estimation of quality measures. However, the corresponding procedure to establish and maintain the marker sets is cumbersome and requires manual curation to ensure the consistency of marker sets [1]. Because the detection of protein domains relies on Profile Hidden Markov Models, for the more comprehensive maker sets, the computational cost noticeably increases when analyzing large metagenomic data sets.

We have developed a new approach for quality assessment which is based on a fast identification of protein domains with UProC [4] and an adaptive clustering of reference genomes. Our method does not depend on a prior identification of marker sets and allows for an easy extension of the database. Adding a new reference genome to the database only requires the computation of the corresponding domain profile with UProC and an automatic redundancy check with respect to the reference profiles that are already contained in the database.

## 2 Methods

The main idea behind BinChecker is to cluster all potential reference genomes in order to identify a suitable subset of genomes that can be used for a protein feature-based estimation of bin completeness and contamination. However, BinChecker does not rely on an a priori clustering of reference genomes. This would result in a partitioning of genomes that is independent of the query profile of the bin to be analyzed. Instead, we use the bin profile to construct a feature space that is used for a query-dependent clustering. Therefore, the final subsets of genomes and features used for estimation are specifically obtained for each individual bin. In the following, we describe all computational steps of our approach in detail.

### 2.1 Pfam protein domain detection

For the identification of marker genes we utilize UProC [4] in combination with protein family database PFAM [3] version 28. We do not use the most recent PFAM version because it significantly increases the UProC memory requirements. In that way, we could successfully run BinChecker on machines with 16 GB of memory. We used UProC with the included ORF detection and the default noise threshold 10*-*3. For each bin and all reference genomes we compute the PFAM profile of all domain counts which give rise to a high-dimensional sparse vector representation.

### 2.2 Database construction

The BinChecker reference database comprises two parts, according to bacterial and archaeal genomes. Both parts are constructed independently. The first step is to calculate the Bray-Curtis dissimilarity between reference profiles, which is half the Manhattan distance between relative frequency vectors. Database construction starts with the two most divergent reference profiles and iteratively adds further reference profiles. For highly similar genomes with a dissimilarity of profiles below 1%, a single representative was selected by chance. This prevents the redundancy that would otherwise arise from very similar strains or genomes that have been re-sequenced multiple times. Also, when adding future reference profiles, this redundancy check would have to be performed. For the construction of the current version of the database we obtained 12744 complete genomes from NCBI including all bacterial and 421 archaeal genomes, published before 1st January 2019. The above similarity filtering resulted in a final number of 9886 reference genomes.

### 2.3 Estimation algorithm

For each bin to be analyzed the following steps are performed.

#### Kingdom classification

First, the representative kingdom for the bin is determined. For this step, we use the Taxy-Pro method [5] with the EM-algorithm, which provides an estimate of the taxonomic profile. This is used to decide if the bin is closely related to bacteria or archaea. The appropriate kingdom is selected if either archaea or bacteria make up more than 80% of the profile. For unclassified bins below that threshold only protein features occurring as SCG in at least 95% of all bacterial and archaeal reference genomes are used for estimation. All later steps of the algorithm are skipped for these bins. The kingdom classification reduces the possible number of reference genomes and therefore increases the speed of the algorithm. In addition, it reduces potential outliers that could arise from an inadequate application of the following more sensitive steps of the algorithm.

#### Feature space transformation and clustering

If we denote the ratio of bin to reference counts for the *i*-th protein feature as 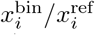 we can calculate specific contamination (cont) and completeness (comp) values from the profile comparison with each reference genome:

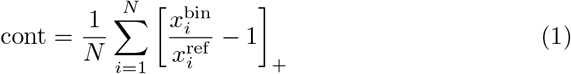

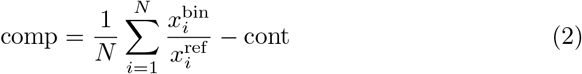

with [*z*]_+_ = max(*z*, 0). Consequently, each reference genome is represented by a point in the comp vs. cont plane and we can use these 2D coordinates for clustering and selection of reference genomes in a drastically reduced feature space. For density-based clustering we use the OPTICS algorithm [6] which is able to detect clusters with different local density. OPTICS requires the specification of an *ϵ* value and a minimum number of points for a cluster. For the number of points, we use the default value 5. To find the optimal value for the *ϵ* radius parameter, the maximum inner cluster distance, we start with a radius of 0 where every point is a singleton cluster. Then we count the decreasing number of clusters for an increasing value of *ϵ* and detect a suitable number of clusters according to an elbow criterion. The initial clusters are then extended by addition of close-by points until each cluster reaches an area with a maximum size of 9*πϵ*. For the extension, BinChecker utilizes the hierarchical structure of the data induced by the OPTICS algorithm. Besides the clustering, we also identify the reference point closest to the origin which we refer to as the first reference point. This point is used as a reference for measuring relative distances in the 2D plane in all further steps.

With regard to the clustering process it now becomes clear that the prior feature transformation to the completeness vs. contamination space is important for multiple reasons: first it provides a huge speed-up of the clustering step. Secondly, it eliminates high-dimensional noise that can significantly deteriorate the clustering. Finally, the dimensionality reduction also provides a meaningful visualization of the clustering results.

#### Marker subset selection

For all references in one cluster, the program identifies a set of marker genes with consistent protein feature counts. The criterion is that at most 5% of the protein features can deviate by more than 10% from the mean count. In general, a cluster with a higher density results in a larger number of selected features. BinChecker combines all obtained feature sets in a final step, trying to successively join the three and five closest clusters to the first reference point. The combination of different sets has to fulfill the same consistency criterion as before, now checking the feature deviations from the overall mean counts of the combined clusters. The number of selected features generally decreases during this aggregation process, which always starts with all features from the first reference point.

### 2.4 Test data generation

In addition to the reference genomes we also obtained complete microbial genomes for testing from NCBI, published 2020 and 2021. These “novel” genomes were used to simulate bins with known completeness and contamination. According to the above similarity filtering we achieved 3987 genomes for testing. Genomes were cut into fragments with an approximate length of 15kb for contig simulation. The completeness of simulated bins was randomly sampled from a range between 60 and 100% while the contamination was between 0 and 40%. Genomes for realistic contamination of a bin were selected from candidate genomes with a high k-mer similarity to the actual (”true”) genome of that bin. In that case, Bray-Curtis similarity of hexamer profiles was between 80 and 100%. If multiple candidates within that similarity range were available, the contaminants were chosen at random. According to this scheme we simulated 10000 bins for a comparative evaluation of BinChecker and CheckM.

## 3 Results

CheckM and BinChecker were applied to the 10000 simulated bins and prediction error was measured by the average absolute deviation from the simulated completeness and contamination values. Table 1 shows the error of both methods together with the standard deviation in brackets. Because we realized some severe outliers with both methods, we eliminated test genomes which resulted in CheckM completeness estimates below 80% for the putatively complete genome sequence. We cannot exclude that these genomes have falsely been classified as ”complete”. The additional filtering slightly reduced the number of test genomes from 3987 to 3921 and from 10000 simulated bins to 9571.

**Table 1.** Average prediction error on test data

| Method | completeness error (± std) | contamination error (± std) |
| --- | --- | --- |
| CheckM | 4.49% (±5.17%) | 5.79% (±5.09%) |
| BinChecker | 4.96% (±7.24%) | 5.33% (±6.18%) |

The results show that BinChecker performance is close to that of CheckM, with a slightly higher error for completeness and a slightly lower error for contamination estimates. Both methods show a considerable standard deviation of the error, with slightly higher values for BinChecker. With regard to overall runtime, Binchecker is about twice as fast as CheckM. The protein feature identification with UProC only requires about 10% of the computation time. Most time consuming parts are the kingdom classification with Taxy-Pro and the OPTICS clustering where each process takes about 40% of the time.

Besides the completeness and contamination estimates, BinChecker also provides an intuitive visualization of the reference genome clustering as additional information for each analyzed bin. An example output is shown in Fig. 1) where the colors indicate the identified clusters of reference genomes in the completeness/contamination-plane. In particular, cluster shape and location with respect to the origin can be meaningful in terms of additional quality indicators.

**Fig. 1.**
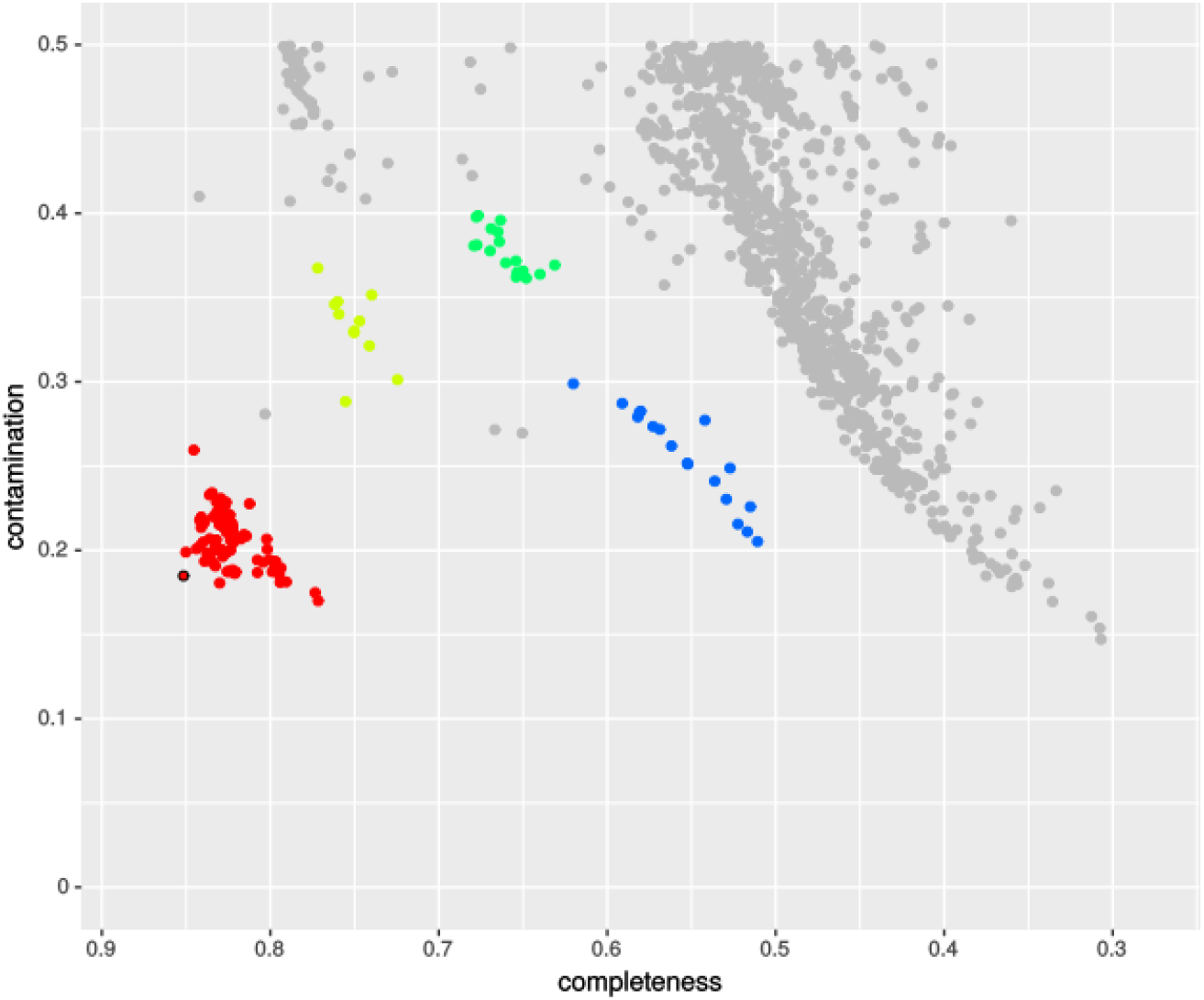
Bin-specific visualization of BinChecker clustering result (see text)

## 4 Conclusion

We have introduced a new algorithm for prediction of bin quality in terms of completeness and contamination of putative genomes. To our knowledge, BinChecker is currently the only alternative to the widely-used CheckM tool. Potentially, BinChecker is able to use more genomic features for estimation because it is not restricted to single copy genes. Since the protein feature sets that are used for estimation are identified for each bin at runtime, there is no training phase for a prior selection of marker sets when building the reference database. In comparison to CheckM, this significantly simplifies the database creation and maintenance. Our simulation results indicate a similar prediction performance and a slightly superior speed of BinChecker with the current implementation. A download link for the Docker container image of the complete BinChecker pipeline is provided at http://gobics.de/BinChecker/

## Acknowledgements

This work was partly funded by Deutsche Forschungsgemeinschaft (DFG, German Research Foundation).

